# Global soil metagenomics reveals ubiquitous yet previously-hidden predominance of *Deltaproteobacteria* in nitrogen-fixing microbiome

**DOI:** 10.1101/2022.12.09.519847

**Authors:** Yoko Masuda, Kazumori Mise, Zhenxing Xu, Zhengcheng Zhang, Yutaka Shiratori, Keishi Senoo, Hideomi Itoh

## Abstract

Biological nitrogen fixation is a fundamental process sustaining all life on earth. In the soil, *Alphaproteobacteria, Betaproteobacteria*, and *Cyanobacteria* are traditionally considered the major groups of nitrogen fixation population. This conventional knowledge has been accumulated by numerous PCR amplicon surveys for nitrogenase genes. However, this prevalent view appears to be inconsistent with results of recent PCR-free meta-omics investigations. Here, we show the global distribution and diversity of terrestrial diazotrophic microbiome, updated by fully leveraging the planetary collections of soil metagenomes along with recently expanded culture collections. After extensive analysis of 1,445 soil metagenomic samples, we proved that the previously-undervalued *Anaeromyxobacteraceae* and *Geobacteraceae* within *Deltaproteobacteria* are ubiquitous and predominant groups of diazotrophic microbiome in the soils, spanning a broad spectrum of environments, with different geographic origins and land usage types, including aerobic (croplands, grasslands) and anaerobic ones (paddy soils, sediments). Our results delineate a revised scheme of soil nitrogen fixation, which is inclusive of both conventional diazotrophs and the so far undervalued *Deltaproteobacteria*.

## Introduction

Biological nitrogen fixation driven by diverse soil microorganisms is a distinct process providing the pedosphere with nitrogen, the major limiting factor for primary production (Vitousek et al., 2002). The identification of microbial players for nitrogen fixation (diazotrophs) in the soil has profound implications for both basic and applied science, since it will improve our understanding of the homeostatic mechanisms taking place in terrestrial ecosystems and will allow the discovery/development of useful microbial resources/technologies for promoting crop production (Soumare et al., 2020). Therefore, since their discovery in the late nineteenth century (Beijerinck, 1888), the distribution and diversity of the diazotrophs in the soil has been one of the most active research topics and is constantly updated along with the accumulation of knowledge and technological innovations (Bürgmann et al., 2004; Hsu and Buckley, 2009; Nelson et al., 2016).

Although nitrogenase genes (*nif*) are conserved in a broad taxonomic range of prokaryotes (Dos Santos et al., 2012), *nif* genes derived from *Alphaproteobacteria, Betaproteobacteria*, and *Cyanobacteria* have been frequently detected in various soil environments such as farmland, grassland, forests, rice paddy fields, riparian zones, and tundra by PCR amplicon surveys targeting *nif* genes (Che et al., 2018; Gaby and Buckley, 2011; Wang et al., 2013; Yu et al., 2018; Zhu et al., 2022). Consequently, these bacteria are considered the primary nitrogen fixers in soil (Kuypers et al., 2018; Mahmud et al., 2020). In contrast, our shotgun metagenomic and metatranscriptomic analyses of paddy soil in Japan have detected highly abundant *nif* genes and transcripts from the families *Anaeromyxobacteraceae* and *Geobacteraceae* within *Deltaproteobacteria* (also classified as the phyla *Myxococcota and Desulfobacterota*, respectively) compared to the findings for the conventional diazotrophic groups (Masuda et al., 2017). Considering their prevalence across many soil types as revealed by 16S rRNA gene-based surveys (Mitter et al., 2021; Pecher et al., 2020; Sun et al., 2018), members of the families *Anaeromyxobacteraceae* and *Geobacteraceae* may thus represent universal and/or major drivers of terrestrial nitrogen fixation. However, these clades, which are well-known iron-reducing bacterial groups (Weber et al., 2006), have received considerably less attention as diazotrophs in soil than the conventional groups because of the less frequent detection of their *nif* genes in soil environments in previous PCR amplicon surveys.

One potential problem is that genomic information of *Anaeromyxobacteraceae* and *Geobacteraceae* has been poorly represented in reference databases because pure isolates of these bacteria have been difficult to obtain. Fortunately, recent studies significantly enriched the reference sequence databases by isolating dozens of novel members within these families using our previously developed slurry incubation method (Itoh et al., 2022, 2021; Liu et al., 2021; Masuda et al., 2020; Xu et al., 2021, 2020, 2019; Yang et al., 2022; Zhang et al., 2021). In addition, it has been reported that diversity analyses based on PCR amplicon sequencing can miss and/or underestimate even major groups in the environmental microbiome because of so-called PCR bias, the poor coverage of target genes by universal primers (Delmont et al., 2018; Jones et al., 2013), calling for extensive analyses of shotgun metagenomic data rather than PCR amplicon-based data. In this context, global-scale shotgun metagenomics is expected to advance current knowledge on the terrestrial diazotrophic microbiota, similarly to the great revisions in diversity fostered in marine environments by recent shotgun metagenomics (Delmont et al., 2018).

In this study, we aimed to reveal the global distribution and diversity of the terrestrial diazotrophic microbiome considering the presence of *Anaeromyxobacteraceae* and *Geobacteraceae* bacteria. We analyzed 1445 shotgun metagenomic datasets to investigate the potential contribution of these clades toward terrestrial nitrogen fixation, making full use of recently published genomic information on *Anaeromyxobacteraceae* and *Geobacteraceae* isolates. Our analyses revealed that *Anaeromyxobacteraceae* and *Geobacteraceae* are the major constituents of the diazotrophic microbiome in terrestrial ecosystems, with their estimated abundance rivaling or exceeding that of the conventional diazotrophic groups.

## Results and discussion

### Diversity of *nif*-harboring genomes in public databases

We first reviewed the currently known diversity of nitrogen-fixing prokaryotes in public databases. KEGG included 7,152 bacterial genomes (KEGG ftp as of August 31, 2022), and 697 of them encoded all three structural genes of nitrogenase, namely *nifH, nifD*, and *nifK*. Among these genomes, those of *Alphaproteobacteria, Firmicutes*, and *Gammaproteobacteria*, as well as *Deltaproteobacteria*, were abundant (Fig. 1a).

**Figure 1.**
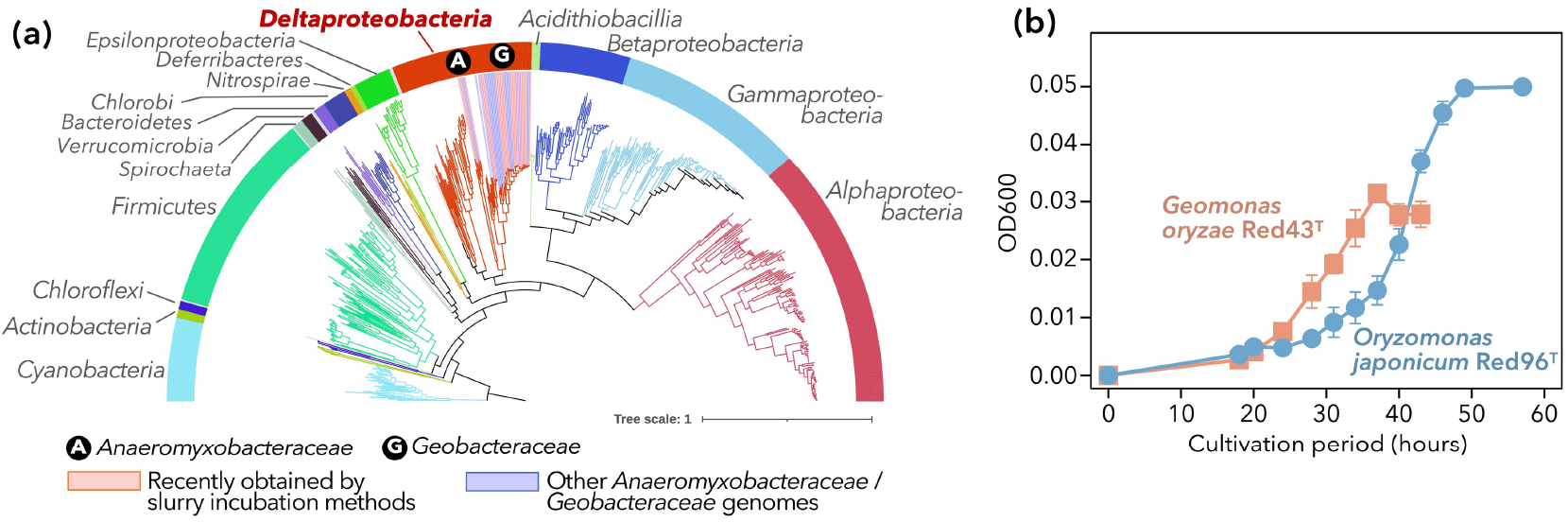
Currently known diversity of diazotrophs. (a) A genome-based phylogenetic tree consisting of potential diazotrophic bacteria [i.e., the genomes of which harbor all three core genes of nitrogenase (*nifH, nifD*, and *nifK*)], including the genomes of new isolates of *Anaeromyxobacteraceae* and *Geobacteraceae* (Table 1). The colors of branches and the band surrounding the tree denote the phyla and proteobacterial classes. Genomes of families *Anaeromyxobacteraceae* and *Geobacteraceae*, the foci of the present study, are highlighted with circled alphabets (A and G) and colored backgrounds (blue and pink, respectively). (b) Growth curves of the type strains of two type species within the family *Geobacteraceae*, namely *Geomonas oryzae* S43^T^ and *Oryzomonas japonica* Red96^T^. The two isolates were grown on MFM medium with N_2_ as the sole nitrogen source. Average and standard deviation of each time point (n = 3) are indicated. Some error bars are shorter than the symbol size.

The representations of *Geobacteraceae* and *Anaeromyxobacteraceae* sequences in public databases were recently improved. At the end of 2018 (prior to the development of the slurry incubation method), RefSeq contained 23 genomes from *Geobacteraceae* and 5 from *Anaeromyxobacteraceae*, whereas these numbers were tripled by September 28, 2022. Approximately 50% of these increases could be attributed to isolates obtained by the slurry incubation method (Table 1) (Itoh et al., 2022, 2021; Liu et al., 2021; Masuda et al., 2020; Xu et al., 2021, 2020, 2019; Yang et al., 2022; Zhang et al., 2021), and all of these isolates bear the core *nif* genes (*nifHDK*) in their genomes. Apart from these isolates, we obtained two other distinct strains belonging to the genus *Geomonas*, namely Red32 (isolated from paddy soil in Joetsu, Niigata, Japan) and Red276 (pond sediment in Myoko, Niigata, Japan; Table 1). These strains displayed 96.3%–97.4% similarity (based on 16S rRNA gene sequences) to all *Geomonas* type strains, which was below the standard threshold (98.65%) for species delineation (Kim et al., 2014). The genomes of these strains also encoded *nifHDK*.

**Table 1.**
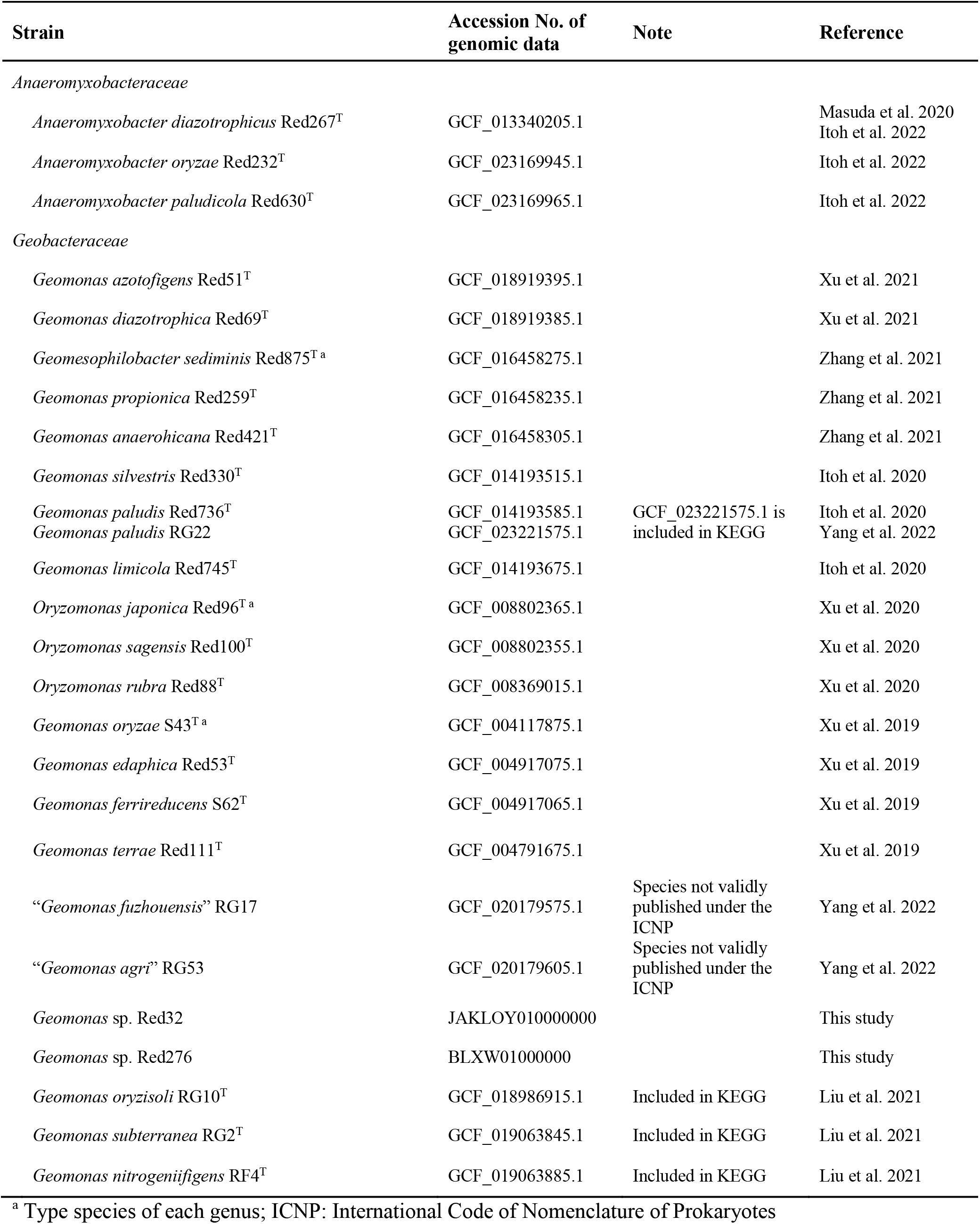
Novel bacterial members within the families *Anaeromyxobacteraceae* and *Geobacteraceae* that have been recently isolated using a slurry incubation method and published to date, including ones isolated in this study. Relevant publications, as well as genomic information, are also indicated.

In addition to the presence of *nif* genes on the genomes, we confirmed the nitrogen-fixing activities of bacterial strains from these clades. In this study, we demonstrated that two type species within *Geobacteraceae*, namely *Geomonas oryzae* S43^T^ and *Oryzomonas japonica* Red96^T^ (Xu et al., 2020, 2019, Table 1), were able to grow on N_2_ as the sole nitrogen source (Fig. 1b). While the acetylene reduction activity of some strains within *Geobacter* and *Geomonas* has been previously tested (Bazylinski et al., 2000; Xu et al., 2021; Liu et al., 2021), the ammonium-independent steady growth of *Geomonas* and *Oryzomonas* suggests that nitrogen fixation is energetically available. The present result, combined with the previously reported N_2_-dependent growth and acetylene reduction activity of *Anaeromyxobacter* and *Geobacter* strains (Bazylinski et al., 2000; Masuda et al., 2020), indicate that *Anaeromyxobacteraceae* and *Geobacteraceae* are likely to be physiologically relevant to nitrogen fixation. We suspected that the use of their genomes as references would yield a better sensitivity in shotgun metagenomic analyses of deltaproteobacterial diazotrophs.

### Global distribution of diazotrophs in terrestrial environments

To assess the global distribution of nitrogen-fixing populations, we collected 1,433 shotgun metagenomic datasets from public databases, namely NCBI SRA and MG-RAST (Katz et al., 2022; Meyer et al., 2019), coming from various environments including cropland soils, forest soils, grassland soils, paddy soils, sediments (including wetlands), and tundra soils (Fig. 2a and Table S1).

**Figure 2.**
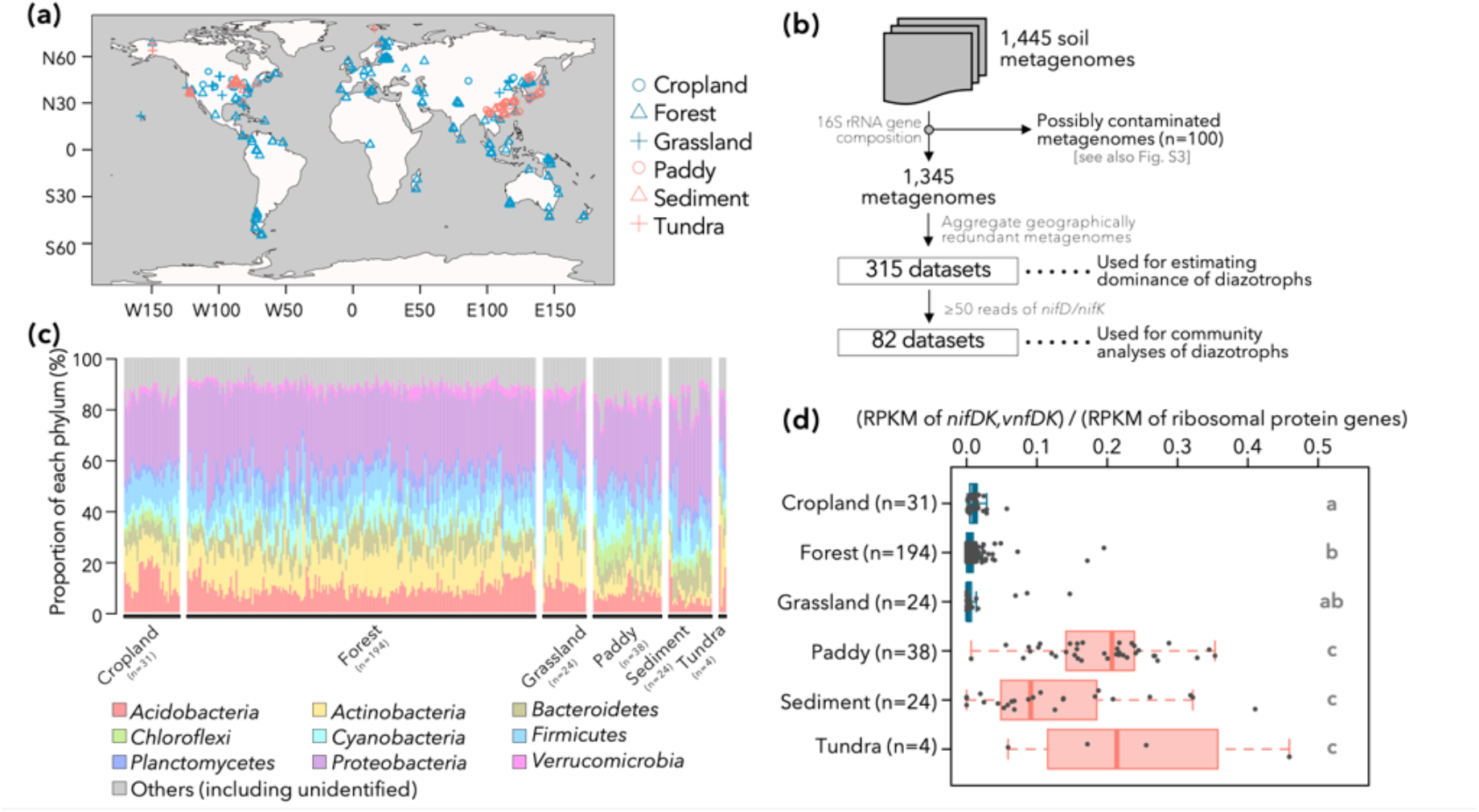
Overview of metagenomic datasets used in this study and distribution of diazotrophs therein. (a) The sampling locations for each metagenomic datasets used in this study. Six types of environments are differentiated by the shapes and colors of symbols. (b) Filtering procedure of metagenomic datasets. The filtering criteria, as well as the aggregation of geographically similar samples, are explained in the panel. (c) Phylum-level prokaryotic community structure of the 315 metagenomic datasets estimated by 16S rRNA gene sequences. (d) The dominance of nitrogen-fixing population in each environment. For each of the 315 metagenomic datasets, the ratio of reads per kilobase of reference sequence per million sample reads (RPKM) of nitrogenase genes to the RPKM of ribosomal protein genes is displayed. The alphabets on the right side of the box indicates the statistical significance in RPKM ratio between different environmental categories (*P* < 0.05, Brunner– Munzel test with Bonferroni’s correction). First, second, and third quantiles are indicated by solid lines. The whiskers, if any, denote 1.5*[interquartile range] from first or third quartile.

Since metagenomes of Japanese soils (including volcanic soils) were poorly represented in public databases, we also collected 12 soil samples in Japan and sequenced their metagenomes. Following preprocessing and curation of these datasets (i.e., merging of paired-end reads, quality filtering), we identified reads bearing nitrogenase genes (*nifD, nifK, vnfD*, and *vnfK*), 16S rRNA genes, and single-copy ribosomal protein genes conserved in most bacterial genomes (Parks et al., 2022) (Table S2). The phylogenetic compositions of 16S rRNA gene sequences indicated that 100 of these datasets could be contaminated by members of order *Lactobacillales* or plants’ plastids (amounting up to 80.2% of *Lactobacillales* or 71.9% of Chloroplast 16S rRNA gene reads: Fig. S1), and the remaining 1,345 were used for further analyses (Fig. 2b). Some of the 1345 metagenomes were redundant and dependent on each other (e.g., technical replicates). Therefore, we combined metagenomes taken from environmental samples within 1 km of each other. The total dataset included 315 samples (Fig. 2b), each considered to bear independent information. The 315 samples were overall dominated by well-known soil-dwelling bacterial clades such as phyla *Proteobacteria, Acidobacteria, Actinobacteria*, etc. (Fig. 2c) and showed no obvious hallmarks of abnormality or technical contamination.

From each of the 315 samples, we detected 122–6,275,556 reads (median: 7,364 reads) of ribosomal protein genes listed in Table S2. We estimated the relative abundances of nitrogen-fixing prokaryotes using the number of nitrogenase gene reads normalized by the number of ribosomal protein gene reads, taking into account the differences in gene lengths between orthologs (corrected by RPKM: see Methods). The relative abundances of nitrogenase gene reads were higher in paddy soils, sediments, and tundra soils (i.e., anaerobic environments) than those in aerobic environments, namely cropland, forest, and grassland soils (*P* < 0.05 in pairwise Brunner–Munzel test with Bonferroni correction, Fig. 2d). On average, nitrogenase genes were detected 17.1 times more frequently in the anaerobic samples than in aerobic samples. This is consistent with both the well-established notion that biological nitrogen fixation is an anaerobic process and the oxygen-sensitive nature of nitrogenase (Robson and Postgate, 1980).

Relative abundances of nitrogenase gene reads exhibited major variations among samples from aerobic environments (i.e., cropland, forest, and grassland), with some harboring low numbers, and others exceptionally dominance of diazotrophs. Cropland and grassland samples may be affected by the roots of leguminous plants and nodule symbionts therein. Some samples from aerobic environments may be locally aerobic, and this might explain the variance in the relative abundance of diazotrophs. We also acknowledge the ambiguity in distinguishing forest or grassland soils from wetland sediments. For example, samples from Disney Wilderness Preserve (DWP), which are labeled as “area of pastureland or hayfields” in MG-RAST and presented the highest relative abundances of nitrogenase genes among “grassland” samples, may have originated from wetland-like environments, as the landscape of DWP bears patches of wetlands (Drake and Weishampel, 2000).

### Global diversity of diazotrophs in terrestrial environments

The taxonomic compositions of diazotrophic communities were further investigated for a more limited dataset of 82 samples, each of which comprised at least 50 sequences of *nifD/K* (Fig. S2ab). Reads encoding *nifD/K* from class *Deltaproteobacteria*, especially *Geobacteraceae* and *Anaeromyxobacteraceae*, were consistently dominant in anaerobic environments such as paddy soils and sediments, as well as in some of the aerobic samples (Fig. 3ab).

**Figure 3.**
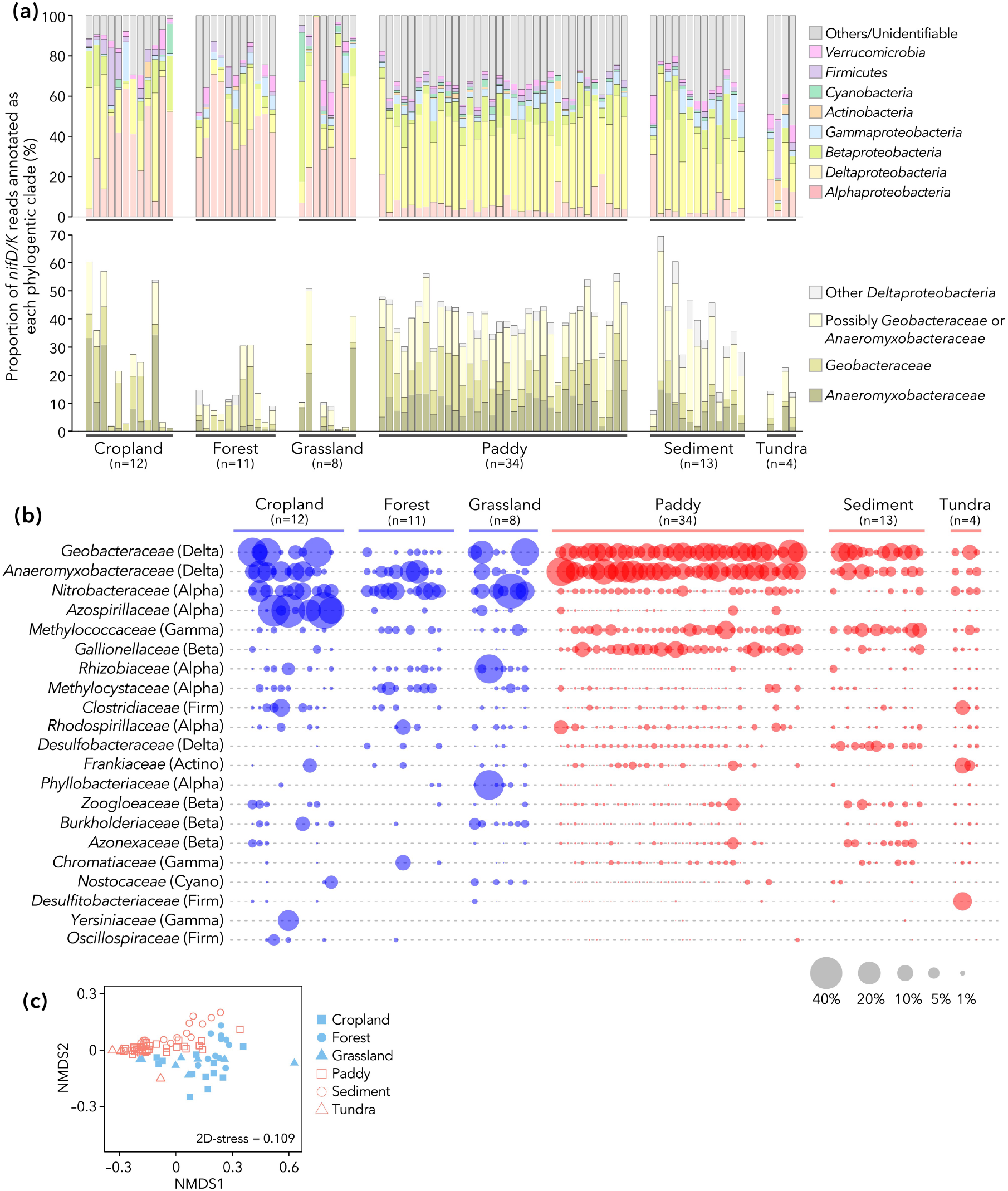
Phylogenetic compositions of nitrogenase genes in the metagenomic datasets with at least 50 reads of *nifD* and *nifK* (n = 82 in total). (a) Upper panel: phylum- and proteobacterial class-level composition. Lower panel: breakdown of deltaproteobacterial composition at the family level. (b) Family-level distribution of *nifD* and *nifK* reads. The correspondence with the phylum- and proteobacterial class-level taxonomy is noted in parentheses: Delta, *Deltaproteobacteria*; Alpha, *Alphaproteobacteria*; Gamma, *Gammaproteobacteria*; Beta, *Betaproteobacteria*; Firm, *Firmicutes*; Actino, *Actinobacteria*; Cyano, *Cyanobacteria*. The area size (not the radius) of each plot is proportional to the proportion of each family within each dataset. (c) The overall beta-diversity of *nifD* and *nifK* sequences summarized by nonmetric multidimensional scaling (NMDS). The shape of each plot denotes the type of environment each dataset comes from. The left panel includes all of the 82 datasets, while the right panel shows NMDS scores calculated after removal of an outlier dataset (one denoted by pink circle in the left panel). The stress values for NMDS are indicated in the panels.

Other major clades within nitrogen-fixing populations included *Alpha*proteobacteria, such as *Nitrobacteraceae* and *Rhizobiaceae* (Fig. 3ab), although some (e.g., *Bradyrhizobium* and *Rhizobium*) in these families are symbiotic diazotrophs and thus possibly incapable of independent nitrogen fixation outside their host plants. The community compositions were significantly different between the aerobic and anaerobic samples (permutational analysis of variance (PERMANOVA) of UniFrac distances, R^2^ = 0.11, *P* = 0.001), as further evidenced by distinct grouping of the two types of samples in nonmetric multidimensional scaling (NMDS) analysis (Fig. 3c). These results indicate that members of *Geobacteraceae* and *Anaeromyxobacteraceae* are the prominent drivers of nitrogen fixation in terrestrial ecosystems. While previous studies in wheat-soybean rotation croplands (Fan et al., 2019) and paddy soils (Masuda et al., 2017; Wang et al., 2021) are in line with our results, the metagenomic datasets analyzed here covering a wide range of environments provide a generalizable insight into the contributions of these clades to nitrogen fixation processes in the pedosphere.

The prevalence of diazotrophic *Geobacteraceae* has actually been reported in some of amplicon sequencing studies (Fan et al., 2019; Wang et al., 2022), but the dominance of *Anaeromyxobacteraceae* has been overlooked in these. This may be due to the high GC content of nitrogenase genes in *Anaeromyxobacteraceae* (65.6%-69.7%, Fig. S3), which may have hindered their amplification by PCR. In congruence with this speculation, 16S rRNA genes of *Anaeromyxobacteraceae*, which have rather moderate GC contents (57.6%–58.6%, Fig. S3) compared to that of their *nif* genes, have been constantly detected by PCR amplicon surveys (Mitter et al., 2021; Pecher et al., 2020; Sun et al., 2018). However, note that GC richness may not solely explain why *Anaeromyxobacteraceae* have been overlooked as diazotrophs in amplicon surveys, since there are examples of high GC content (63.4%–68.0%) nitrogenase genes belonging to *Frankia* species (phylum *Actinobacteria*) that are successfully amplified by PCR-based methods (Collavino et al., 2014; Dahal et al., 2017).

Another characteristic of the diazotrophic communities of anaerobic environments (with the exception of tundra) is the high similarity between samples (Fig. 3). While all the paddy soil and sediment samples are dominated by *Deltaproteobacteria* and presented overall low beta-diversity levels, environments such as cropland, forest, and grassland are dominated by more diverse clades of diazotrophs showing high beta-diversity levels. One possible explanation is the heterogeneity among aerobic samples: some of the cropland, forest, and grassland samples may be associated with leguminous vegetations (i.e., affected by nodule-associated bacteria) or originated from locally muddy soils, while others are not. Another explanation for this divergence in community structures is ecological drift (Nemergut et al., 2013; Vellend, 2010). Diazotrophs have smaller population sizes in aerobic samples (Fig. 2d); thus, their communities are expected to be more sensible to ecological drift, resulting in increased beta-diversities between communities as previously shown (Fodelianakis et al., 2021).

Although we used a limited dataset of 82 samples bearing at least 50 *nifDK* sequences for the taxonomic composition analysis, the bias introduced by this manipulation is unlikely to be critical. First, the selected samples do not necessarily present high relative abundances of diazotrophs, since the number of total reads greatly varies between samples (Fig. S2ab). Second, the abundance ratio of nitrogenase genes to ribosomal protein genes explain only 5.5% the phylogenetic diversity of *nifDK* (R^2^ = 0.055, *P* = 0.028) (Fig. S2c).

### An approximate estimation of global dominance of deltaproteobacterial diazotrophs

Analyses of the global metagenomic dataset indicated that anaerobic environments harbor high abundances of diazotrophic prokaryotes and *Anaeromyxobacteraceae* and *Geobacteraceae* are the dominant diazotrophs in these environments. Wetlands (possibly including waterlogged paddy soils) represent between 5.2% (Xu et al., 2013) and 8% (Davidson et al., 2018) of all lands, with microbial biomass carbon therein amounting to 10.3% of the microbial biomass in all lands [calculated from the data presented in (Xu et al., 2013)]. However, given that the relative abundance of nitrogen fixers was 17.1 times higher in anaerobic microbial communities than in aerobic ones (Fig. 2d), anaerobic soils are expected to harbor a large proportion of soil nitrogen fixers. This further suggests that *Anaeromyxobacteraceae* and *Geobacteraceae*, nitrogenase genes of which show higher relative abundance in anaerobic soils (Fig. 3b), are the predominant diazotrophs at the global terrestrial ecosystem scale. Unfortunately, it is impossible to quantitate the contribution of *Anaeromyxobacteraceae* and *Geobacteraceae* to nitrogen fixation in soil environments based on the results of this study alone, since physiological traits are not always coupled with presence/absence of genes. However, our results strongly suggest that *Anaeromyxobacteraceae* and *Geobacteraceae* are important diazotrophic members in soil microbial communities worldwide.

It should be noted that the abundance of diazotrophic *Anaeromyxobacteraceae* may be underestimated even in shotgun metagenomic sequencing. Illumina sequencing technology is known to be biased against sequencing GC-rich nucleotide fragments even in shotgun sequencing (Sevim et al., 2019). Considering this bias, the proportion of *Anaeromyxobacteraceae* in the soil, the nitrogenase genes of which are GC-rich (66.9%–69.0%, 65.6%–67.0%, and 66.9%–69.7% for *nifH, nifD*, and *nifK*, respectively, Fig. S3), might be even higher than estimated in this study. Long-read sequencers (i.e., PacBio and Nanopore) are less prone to GC bias (Sevim et al., 2019) and may potentially rectify this issue, although their current yield is orders of magnitude smaller than those of short-read sequencers such as Illumina HiSeq and NovaSeq, and thus currently not a good fit for the characterization of samples as heterogenous and rich in diversity as soil metagenomes.

### Benefits of expanding culture collection

Previous and current efforts to enrich culture collections (Itoh et al., 2022, 2021; Masuda et al., 2020; Xu et al., 2021, 2020, 2019; Zhang et al., 2021) has substantially expanded the available repertoire of *Anaeromyxobacteraceae* and *Geobacteraceae* strains. In fact, an average of 55.4% and 25.6% of NifD/K sequences derived from *Anaeromyxobacteraceae* and *Geobacteraceae* members, respectively, displayed higher proximity to our novel strains than to any other nitrogenase sequence in KEGG from these families (Fig. 4).

**Figure 4.**
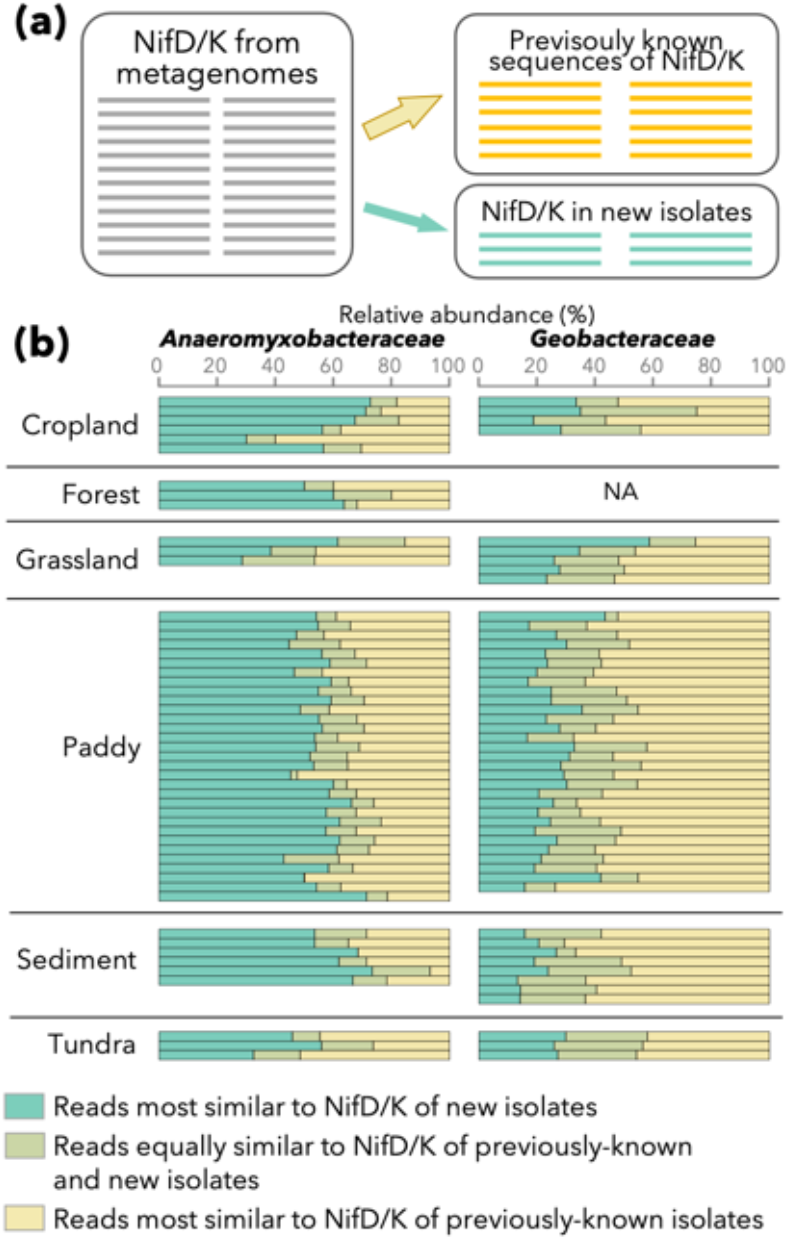
Contribution of nitrogenase gene sequences from newly isolated strains in the bioinformatic analyses of metagenomes. (a) A schematic of the analysis. NifD/K sequences annotated as *Anaeromyxobacteraceae* or *Geobacteraceae* in metagenomes were mapped onto already known sequences of NifD/K (right-upper) and those in our new isolates (right-bottom). Only the top hit for each query sequence (i.e., one from metagenomes) was considered. (b) Relative abundance of metagenome-derived NifD/K sequences that were most similar to already known sequences (yellow) and those from our new isolates (green), as well as those equally similar to the nitrogenase genes of already known genomes and our new isolates (dim green), are summarized. Only datasets with 10 or more sequences of NifD/K for each family are displayed. The proportion of metagenome-derived NifD/K sequences that were most similar to already known sequences is shown in yellow, to sequences from the new isolates in green. Dim green is used to show the proportion of metagenome-derived NifD/K sequences equally similar to nitrogenase genes of already known genomes and of new isolates. Only datasets with ten or more sequences of NifD/K for each family are displayed.

Interestingly, this trend was consistent among a wide variety of environments including aerobic and anaerobic environments, although the majority of the novel strains were isolated from paddy soils or sediments under anaerobic conditions. Based on the present and previous findings, paddy soils and sediments appear to be promising environments for isolating free-living diazotrophs, representing diverse terrestrial environments including aerobic environments such as cropland, forests and grassland.

### Conclusions and outlook

Contrary to the conventional view, our large-scale comparative metagenomics analyses revealed the global distribution and substantial abundance of *Anaeromyxobacteraceae* and *Geobacteraceae* in terrestrial diazotrophic microbiome, highlighting the importance of *Deltaproteobacteria* members (phyla *Myxococcota* and *Desulfobacterota*) in terrestrial ecosystems. Moreover, thousands of obtained nitrogenase sequences exhibited high proximity to our newly isolated strains. Our results warrant further efforts to enrich the culture collections, which would fill the knowledge gaps in the diversity and ecology of diazotrophs. Furthermore, knowledge on terrestrial diazotrophic microbiota and strategies for their cultivation and isolation might be also updated, for example, by using single-cell sorting. In addition, the insights into the contribution of each diazotrophic taxon to nitrogen fixation will be foreseeable, for instance, through stable isotope probing with ^15^N_2_ under more natural conditions, using various soil samples.

## Materials and Methods

### Isolation and genomic sequencing of new soil strains

The *Geomonas* strains Red32 and Red276 were isolated from paddy soil (Joetsu, Niigata, Japan) and pond sediment (Myoko, Niigata, Japan) following the slurry incubation method used to isolate new members of *Geobacteraceae* (Xu et al., 2020, 2019). The soils collected from the paddy field in Nagaoka, Niigata, Japan were air-dried, placed in a 15-mL serum bottle and suspended in distilled water (soil:water, 2:3, *w*/*v*). After autoclaving at 120°C for 20 min, 0.1 g of undried soil were added to the bottle as a microbial inoculum with and without vitamin solution for strains Red276 and Red32, respectively (Masuda et al., 2020). Then, we sealed the bottles with butyl rubber stoppers and aluminum caps, replaced headspace gas with with N_2_/CO_2_ (80:20, *v*/*v*), and incubated them at 30°C for 2 weeks without shaking. Afterward, 200 μL of incubated soil slurry was transferred to the fresh soil slurry bottle and incubated at 30 °C for 2 weeks. After repeating this step once (for strain Red276) or twice (for strain Red32), the incubated soil slurry was streaked on 1.5% agar plates of the R2A broth “DAIGO” (Nihon Pharmaceutical, Tokyo, Japan) supplemented with 5 mM disodium fumarate. The plates were incubated at 30°C for 10 days under anaerobic conditions using the AnaeroPack system (Mitsubishi Gas Chemical, Tokyo, Japan). Red-colored colonies, a typical hallmark of *Geobacteriaceae* strains (Coates et al., 1996; Itoh et al., 2021; Lovley et al., 1993; Xu et al., 2020, 2019), were purified by a single-colony isolation using the same medium plates. Genomic DNA was extracted from the two isolated strains using a DNeasy Blood and Tissue Kit (Qiagen, Hilden, Germany) and sequenced using an Illumina HiSeq sequencer (Illumina, CA, USA). Subsequently, the drafted genomes were generated as previously described (Xu et al., 2020, 2019).

### Diazotrophic activity assay

Following a 5-day culture in nitrogen-free modified freshwater medium (MFM) as previously described (Xu et al., 2020, 2019), the cells of *Geomonas oryzae* S43^T^ and *Oryzomonas japonica* Red96^T^ were transferred to serum bottles containing 20 mL of nitrogen-free MFM (Xu et al., 2019). Bacterial growth was monitored by measuring the suspension absorbance using a spectrophotometer (UV-1900 UV-visible spectrophotometer, Shimadzu, Kyoto, Japan) at a wavelength of 600 nm. The experiments were performed in triplicate.

### Preparation of custom database

We used KEGG database (as of August 31, 2022) for functional gene annotations (Kanehisa et al., 2021). To increase the sensitivity for genes from *Anaeromyxobacteraceae* and *Geobacteraceae*, we customized KEGG database by adding genomes belonging to these families obtained using slurry incubation methods (Table 1). Genomes already included in KEGG were not added. We predicted their coding sequences (CDS) using Prodigal (Hyatt et al., 2010) with default parameters, annotated them using KofamScan version 1.3.0 and KOfam version 2022-08-01 with default parameters (Aramaki et al., 2020), and concatenated the CDS with KEGG database (including those received no K number).

### Phylogenetic analysis of bacterial genomes harboring *nif* genes

From the aforementioned custom database, we screened genomes harboring a set of *nif* core genes, namely *nifH* (K02588 in KEGG), *nifD* (K02586), and *nifK* (K02591). Archaeal genomes were excluded from analysis. The universal single-copy gene sequences were identified from each genome, translated amino acid sequences, and mapped onto multiple sequence alignment (MSA) of GTDB R207 using GTDB-Tk v2.1.0 (Chaumeil et al., 2019; Eddy, 2011; Hyatt et al., 2010; Parks et al., 2022). Here “identify” and “align” commands were used with default parameter settings. The MSA was fed into FastTree 2.1.11 (default parameters) and a phylogenetic tree was constructed (Price et al., 2010). The tree was manually rerooted using *Cyanobacteria* as the outgroup (Hug et al., 2016) and visualized on iTOL server (Letunic and Bork, 2021).

### GC content of 16S rRNA genes and nitrogenase genes among bacterial genomes

Ribosomal RNA genes were identified from each of the bacterial genomes in the custom database explained above using barrnap version 0.9 (https://github.com/tseemann/barrnap; accessed March 22, 2022). Only 16S rRNA sequences with 1000 bases or longer were picked. For each genome with at least one valid 16S rRNA gene sequence and all of the identified *nifH, nifD* and *nifK* (identified as previously described), the GC contents of 16S rRNA genes, *nifH, nifD*, and *nifK* were calculated. When a genome had multiple copies of each gene, GC content was calculated for the concatenated sequence of these copies. Any ambiguous base was excluded from calculation of GC content.

### Soil collection and metagenomic sequencing

We collected 12 soil samples from various agricultural fields in Japan at a depth of 0–5 cm (Table S1). Following the removal of plant residues and additional water from the surface, the soil samples were transported to the laboratory and stored at −80°C until further use for DNA extraction. Soil DNA was extracted from 0.5 g (wet weight) of each soil sample using the ISOIL for Beads Beating Kit (Nippon Gene, Tokyo, Japan) according to the manufacture’s instruction with the following modifications: prior to the beads beating step, 0.02 g skim milk was added to the lysis buffer to improve the extraction efficiency (Takada-Hoshino and Matsumoto, 2004) and post-elution purification using RNase A (Takara, Shiga, Japan) and DNA Clean & Concentrator (Zymo Research) according to the manufacture’s introduction. Purified DNA was quantified using Qubit 2.0 Fluorometer (Invitrogen, Carlsbad, CA, USA) with Qubit dsDNA HS Assay Kits (Invitrogen). The construction of DNA libraries, shotgun sequencing on an Illumina MiSeq sequencer, and merging of paired-end sequences were performed as described previously (Masuda et al., 2017).

### Collection of publicly available metagenomic data and their quality assessment

We further collected reusable bulk soil metagenomic datasets on INSDC (Arita et al., 2021) and MG-RAST (Meyer et al., 2008) that met the following criteria: (i) derived from outdoor samples exempted from post-sampling treatments that can affect the microbial community structure; (ii) sequenced on Illumina MiSeq, HiSeq, MiniSeq, NextSeq or NovaSeq (i.e., state-of-the-art, highly accurate sequencers); and (iii) reported in peer-reviewed literature (with the exception of data obtained by the National Ecological Observatory Network). Moreover, the datasets from rhizosphere soils were not used in this study because they are extensively and dynamically affected by the plant roots (Zhalnina et al., 2018) and not representative of the soil microbial communities. In total, we collected 1445 datasets as listed in Table S1 (Angle et al., 2017; Bahram et al., 2018; Berkelmann et al., 2020; Black and Just, 2018; Cania et al., 2019; Cha et al., 2021; Chen et al., 2019; Chu et al., 2018; Crits-Christoph et al., 2018; Hartman et al., 2017; Huber et al., 2018; Jiang et al., 2018; Johnston et al., 2016; Li et al., 2018, 2020; Links et al., 2021; Liu et al., 2018; Ma et al., 2018; Neal et al., 2021; Nelkner et al., 2019; Orellana et al., 2018; Ouyang and Norton, 2020; Paungfoo-Lonhienne et al., 2017; Romanowicz et al., 2021; Sukhum et al., 2021; Suttner et al., 2020; Hang Wang et al., 2021; Wang et al., 2020; Woodcroft et al., 2018; Wu et al., 2021; Xiao et al., 2016; Xue et al., 2019; Yu et al., 2019; Yurgel et al., 2019; Zhang et al., 2019; Zheng et al., 2021). The latitude and longitude of each sampling site were obtained from public databases [INSDC BioSamples database (Courtot et al., 2022) and MG-RAST] and verified with the descriptions in each publication. The NCBI SRA Toolkit and MG-RAST API (with the option “file=050.1”) were used to transfer the massive data to a domestic supercomputer.

The paired-end sequences were merged using the “fastq_mergepairs” command in USEARCH v11.0.667 (Edgar, 2010) with the options “-fastq_maxdiffs 5 -fastq_minovlen 15”. As a rule, merging of paired-end reads is strongly influenced by the length of the Illumina library, which differs widely between research projects. In particular, read pairs from long libraries (e.g., those designed for metagenomic assembly) are rarely merged. Therefore, we used Read 1 sequences that failed to be merged in addition to successfully merged sequences and single-end sequences. From each of the merged and unmerged sequences, we picked a partial sequence with an expected number of errors of fewer than 0.5 bases. From these partial sequences, those shorter than 150 bases were discarded; therefore, the expected error rate of each sequence was approximately 0.33% (i.e., 0.5 bases per 150 bases) or lower.

To determine the prokaryotic community structure of each dataset, we identified 16S rRNA gene sequences from these filtered sequences, using SortMeRNA v2.1 (Kopylova et al., 2012) with SILVA v138.1_SSURef_NR99 (Quast et al., 2012) as the reference database. Retrieved 16S rRNA gene sequences were further filtered by BLASTn search against SILVA at the maximum e-value threshold of 1e–10 and a minimum query coverage of 50%. Those passing this filter were taxonomically annotated using SINTAX (Edgar, 2016) implemented in USEARCH v11.0.667 with a minimum confidence threshold of 0.5. Here again SILVA was used as the reference database, with the following modifications to maximize the taxonomic annotation accuracy: SILVA taxonomy was replaced with NCBI taxonomy (referred to taxmap_embl-ebi_ena_ssu_ref_nr99_138.1), and sequences without valid taxonomic assignment at phylum, class, or order level (typically metagenome-derived sequences) were eliminated. However, organelle sequences (i.e., sequences annotated as mitochondrial and chloroplast according to SILVA taxonomy) were exceptionally retained.

We excluded 100 metagenomic datasets, with a proportion of *Lactobacillales* or Chloroplast 20% or higher (Fig. S1), from downstream analyses because they were considered possibly contaminated. The remaining 1333 metagenomes from public databases and newly sequenced 12 metagenomes, including those from samples taken within < 1 km, were merged and treated as one sample. The distances between samples were calculated based on the latitude and longitude of each sample using geodesic module in GeoPy (https://geopy.readthedocs.io/en/stable/#; accessed Aug 27, 2022).

### Gene annotations of metagenomic reads

To determine the nitrogen-fixing populations within each metagenomic dataset, the filtered sequences were subjected to homology search against the custom database explained above (i.e., KEGG database supplemented with *Anaeromyxobacteraceae* and *Geobacteraceae* genomes). This search was a two-step process (Yu and Zhang, 2013). First, we mapped the query sequences against nitrogenase amino acid sequences [K02586 (NifD), K02591 (NifK), K22896 (VnfD) and K22897 (VnfK)] using the blastx command implemented in DIAMOND v2.0.9.147 (Buchfink et al., 2021) with the options “--sensitive -e 10e-5”. NifH was not used in because of its primary structure, which can be confused with those of other proteins irrelevant to nitrogen fixation (Mise et al., 2021). Query sequences aligned with nitrogenase sequences, which may be regarded as the candidate *nif* genes, were subjected to a second screening. Specifically, CDSs were inferred from these candidates using Prodigal (metagenomic mode) and mapped against the whole KEGG database to determine whether they were *nif* genes. Once again, the blastp command implemented in DIAMOND was used with the options “--sensitive -evalue 10e-5”. To avoid possible errors caused by the heuristic nature of DIAMOND searches, we obtained up to 200 hits for each query and determined one top hit among them according to the bit scores. Then, a query with a top hit against a K02586 [K02591, K22896 or K22897] sequence was regarded as a *nifD* [*nifK, vnfD* or *vnfK*, respectively] sequence. Note that the top hit was used only to differentiate *nif* genes from other genes, rather than the taxonomic assignment of each query.

To normalize the abundance of the nitrogen-fixing population by the size of the prokaryotic population, we counted the number of reads bearing prokaryotic ribosomal protein genes, single-copy markers conserved among most prokaryotes known to date (Hug et al., 2016). The genes used for this purpose are listed in Table S2, and the reads bearing these genes were identified in the same way as the *nif* genes. Because the length of each marker gene was different (Table S2), we calculated the reads per kilobase of reference sequence per million sample reads (RPKM) for each ribosomal protein gene and used their median as the representative RPKM of the dataset. The length of reference sequence was substituted with the triple of the average length of amino acid sequences in KEGG. We also calculated the RPKMs of four *nif* genes (*nifD, nifK, vnfD* and *vnfK*) to compare them with those of ribosomal proteins. The sum of the four RPKMs was regarded as the “total” RPKM of the *nif* genes. Then we calculated the ratio of the “total” RPKM of the *nif* genes and the median RPKM of the ribosomal protein genes, which represent the population of diazotrophs and total prokaryotes, respectively. This ratio roughly represents double the proportion of diazotrophs to all prokaryotes, as most diazotrophs carry the gene pair “*nifD* and *nifK*” or “*vnfD* and *vnfK*.” For any pair of the environmental categories, the Brunner–Munzel test was performed to test the null hypothesis that RPKM ratios are the same between the two categories.

### Taxonomic assignments of metagenomic reads encoding *nif* genes

The taxonomic annotation of protein-coding genes is a cumbersome and error-prone process because of the rapid evolution of sequences. To make most of the fragmented metagenomic sequences while averting annotation errors, we employed phylogenetic placement, which is implemented in pplacer (Matsen et al., 2010), rather than a simple homology search. The effectiveness of introducing phylogenetic information into taxonomic assignment was previously demonstrated (Kapili and Dekas, 2021). Briefly, an MSA of reference NifD (or NifK) sequences as well as a backbone phylogenetic tree of NifD (or NifK) was constructed. The metagenomic sequences of NifD (or NifK) were mapped onto the backbone tree and were taxonomically annotated. First, NifD (or NifK) sequences in KEGG [under the K numbers K02586 (or K02591)] and our isolates (listed in Table 1) were fed into MAFFT v7.475 (with the option “--auto”) (Katoh, 2002). In this analysis, the identical sequences of NifD or NifK were dereplicated because pplacer’s implementation is incompatible with the zero branch length. None of the identical sequences were derived from different families with only one exception; therefore, this manipulation would not compromise the rigor in taxonomic annotation. From the MSA, an approximate maximum likelihood tree was constructed using FastTree (Price et al., 2009). The tree was automatically rerooted using taxtastic v0.9.2 (distributed along with pplacer) and subsequently used as a backbone phylogenetic tree for phylogenetic placement.

Each NifD/K-like amino acid sequence from the metagenomes was mapped onto the MSA using MAFFT with the option “--add”. That sequence was placed onto the backbone tree using pplacer v1.1.alpha19-0-g807f6f3 (with the option “--mrca-class”), and its phylogenetic taxonomy was determined using guppy v1.1.alpha13-0-g1ec7786 (https://matsen.github.io/pplacer/generated_rst/guppy.html) (with the option “classify --mrca-class”). pplacer and guppy report several or dozens of solutions (i.e., taxonomic annotations) with likelihood values, which enabled us to assess the reliability of each annotation. We used family-, order-, class-, and phylum-level annotations with a likelihood value of 0.5 or higher (the threshold was precisely set at 0.49999 to consider the effect of floating-point errors). Theoretically, two different solutions could satisfy this criterion (with the likelihood values of 0.5 each) for one query, but practically, such a situation did not occur. When none of the annotations satisfied this criterion, the sequence in question was determined as not bearing sufficient information to determine taxonomy. Because the reference sequences for *vnf* (or Vnf) were rather scarce and the reliable taxonomic annotation was not affordable, we did not perform taxonomic annotation of *vnfD/K* sequences. Taxonomic names were handled using TaxonKit (Shen and Ren, 2021) and NCBI Taxonomy System (Federhen, 2012).

### Beta-diversity analyses

Beta-diversity between any pair of diazotrophic communities was calculated between using the average of UniFrac distances for NifD and NifK, which were determined based on the results of phylogenetic placement. We used NMDS with two dimensions to summarize overall beta-diversity between communities. We also performed PERMANOVA with 999 times permutation to test the null hypothesis that community structures of diazotrophs are similar between aerobic and anaerobic environments.

### Homology analyses between NifD/K of metagenomes and isolate genomes

We further mapped the NifD/K sequences annotated as family *Anaeromyxobacteraceae* or *Geobacteraceae* in metagenomic reads to NifD/K sequences from that family in our custom database using the Needleman–Wunsch algorithm implemented in USEARCH. We obtained the sequence similarity between each read and its nearest sequence in the database. We counted the number of reads for which the nearest sequence is from the genomes of bacterial isolates obtained via the slurry incubation method (Table 1).

Throughout the study, we used SeqKit v0.16.1/v2.2.0 (Shen et al., 2016) and R 4.0.5 (R Core Team, 2021), including the package “vegan” (Oksanen et al., 2020), to handle fastq and fasta files and to perform statistical tests, respectively.

## Data availability

Genomic and metagenomic sequences obtained in this study have been deposited in GenBank and DDBJ DRA, respectively. See Tables 1 and S1 for accession numbers.

## Acknowledgements

This work was financially supported by the CANON Foundation, JSPS KAKENHI Grant Numbers JP20H00409, JP20H05679, JP20K15423, and JP22K18029, MEXT KAKENHI Grant Number JP22H04894, JST-Mirai Program grant JPMJMI20E5, and JPNP18016 commissioned by the New Energy and Industrial Technology Development Organization (NEDO). Part of the metagenomic data used in this study was provided by the National Ecological Observatory Network (NEON) via MG-RAST. We thank Haruka Ooi and Emiko Kobayashi (National Institute of Advanced Industrial Science and Technology) for literature survey, Yumi Sugisawa (National Institute of Advanced Industrial Science and Technology) for performing experiments, and anonymous farmers for providing soil samples. Computations were partially performed on the NIG supercomputer at ROIS National Institute of Genetics and the SHIROKANE supercomputer at Human Genome Center, The Institute of Medical Science, The University of Tokyo.

## Author contributions

Y.M., K.M., K.S., and H.I. designed the study and supervised the project. H.I. isolated bacterial strains and Z.X. sequenced their genomes. Z.Z. and Z.X. performed diazotrophy assays. Y.S. and H.I. collected Japanese soil samples, and Y.M. performed metagenomic sequencing of these samples. Y.M. and H.I. performed the primary bioinformatic analysis of the metagenomic dataset. K.M. curated and analyzed genomic and metagenomic datasets. Y.M., K.M., and H.I. wrote the paper with substantial input from all authors.

## References

Angle, J.C., Morin, T.H., Solden, L.M., Narrowe, A.B., Smith, G.J., Borton, M.A., Rey-Sanchez, C., Daly, R.A., Mirfenderesgi, G., Hoyt, D.W., Riley, W.J., Miller, C.S., Bohrer, G., Wrighton, K.C., 2017. Methanogenesis in oxygenated soils is a substantial fraction of wetland methane emissions. Nature Communications 8, 1567. doi:10.1038/s41467-017-01753-4

Aramaki, T., Blanc-Mathieu, R., Endo, H., Ohkubo, K., Kanehisa, M., Goto, S., Ogata, H., 2020. KofamKOALA: KEGG Ortholog assignment based on profile HMM and adaptive score threshold. Bioinformatics 36, 2251–2252. doi:10.1093/bioinformatics/btz859

Arita, M., Karsch-Mizrachi, I., Cochrane, G., 2021. The international nucleotide sequence database collaboration. Nucleic Acids Research 49, D121–D124. doi:10.1093/nar/gkaa967

Bahram, M., Hildebrand, F., Forslund, S.K., Anderson, J.L., Soudzilovskaia, N.A., Bodegom, P.M., Bengtsson-Palme, J., Anslan, S., Coelho, L.P., Harend, H., Huerta-Cepas, J., Medema, M.H., Maltz, M.R., Mundra, S., Olsson, P.A., Pent, M., Põlme, S., Sunagawa, S., Ryberg, M., Tedersoo, L., Bork, P., 2018. Structure and function of the global topsoil microbiome. Nature 560, 233–237. doi:10.1038/s41586-018-0386-6

Bazylinski, D.A., Dean, A.J., Schuler, D., Phillips, E.J.P., Lovley, D.R., 2000. N2-dependent growth and nitrogenase activity in the metal-metabolizing bacteria, Geobacter and Magnetospirillum species. Environmental Microbiology 2, 266–273. doi:10.1046/j.1462-2920.2000.00096.x

Beijerinck, M.W., 1888. Die bacterien der papilionaceenknöllchen. Botanische Zeitung 46, 725–735.

Berkelmann, D., Schneider, D., Meryandini, A., Daniel, R., 2020. Unravelling the effects of tropical land use conversion on the soil microbiome. Environmental Microbiome 15, 5. doi:10.1186/s40793-020-0353-3

Black, E.M., Just, C.L., 2018. The Genomic Potentials of NOB and Comammox Nitrospira in River Sediment Are Impacted by Native Freshwater Mussels. Frontiers in Microbiology 9, 2061. doi:10.3389/fmicb.2018.02061

Buchfink, B., Reuter, K., Drost, H.-G., 2021. Sensitive protein alignments at tree-of-life scale using DIAMOND. Nature Methods 18, 366–368. doi:10.1038/s41592-021-01101-x

Bürgmann, H., Widmer, F., Von Sigler, W., Zeyer, J., 2004. New Molecular Screening Tools for Analysis of Free-Living Diazotrophs in Soil. Applied and Environmental Microbiology 70, 240–247. doi:10.1128/AEM.70.1.240-247.2004

Cania, B., Vestergaard, G., Krauss, M., Fliessbach, A., Schloter, M., Schulz, S., 2019. A long-term field experiment demonstrates the influence of tillage on the bacterial potential to produce soil structure-stabilizing agents such as exopolysaccharides and lipopolysaccharides. Environmental Microbiome 14, 1. doi:10.1186/s40793-019-0341-7

Cha, G., Meinhardt, K.A., Orellana, L.H., Hatt, J.K., Pannu, M.W., Stahl, D.A., Konstantinidis, K.T., 2021. The influence of alfalfa-switchgrass intercropping on microbial community structure and function. Environmental Microbiology 23, 6828–6843. doi:10.1111/1462-2920.15785

Chaumeil, P.-A., Mussig, A.J., Hugenholtz, P., Parks, D.H., 2019. GTDB-Tk: a toolkit to classify genomes with the Genome Taxonomy Database. Bioinformatics 36, 1925–1927. doi:10.1093/bioinformatics/btz848

Che, R., Deng, Y., Wang, F., Wang, W., Xu, Z., Hao, Y., Xue, K., Zhang, B., Tang, L., Zhou, H., Cui, X., 2018. Autotrophic and symbiotic diazotrophs dominate nitrogen-fixing communities in Tibetan grassland soils. Science of The Total Environment 639, 997–1006. doi:10.1016/j.scitotenv.2018.05.238

Chen, Y.-P., Liaw, L.-L., Kuo, J.-T., Wu, H.-T., Wang, G.-H., Chen, X.-Q., Tsai, C.-F., Young, C.-C., 2019. Evaluation of synthetic gene encoding α-galactosidase through metagenomic sequencing of paddy soil. Journal of Bioscience and Bioengineering 128, 274–282. doi:10.1016/j.jbiosc.2019.03.006

Chu, B.T.T., Petrovich, M.L., Chaudhary, A., Wright, D., Murphy, B., Wells, G., Poretsky, R., 2018. Metagenomics Reveals the Impact of Wastewater Treatment Plants on the Dispersal of Microorganisms and Genes in Aquatic Sediments. Applied and Environmental Microbiology 84. doi:10.1128/AEM.02168-17

Coates, J.D., Phillips, E.J., Lonergan, D.J., Jenter, H., Lovley, D.R., 1996. Isolation of Geobacter species from diverse sedimentary environments. Applied and Environmental Microbiology 62, 1531–1536. doi:10.1128/aem.62.5.1531-1536.1996

Collavino, M.M., Tripp, H.J., Frank, I.E., Vidoz, M.L., Calderoli, P.A., Donato, M., Zehr, J.P., Aguilar, O.M., 2014. nifH pyrosequencing reveals the potential for location-specific soil chemistry to influence N_2_-fixing community dynamics. Environmental Microbiology 16, 3211–3223. doi:10.1111/1462-2920.12423

Courtot, M., Gupta, D., Liyanage, I., Xu, F., Burdett, T., 2022. BioSamples database: FAIRer samples metadata to accelerate research data management. Nucleic Acids Research 50, D1500–D1507. doi:10.1093/nar/gkab1046

Crits-Christoph, A., Diamond, S., Butterfield, C.N., Thomas, B.C., Banfield, J.F., 2018. Novel soil bacteria possess diverse genes for secondary metabolite biosynthesis. Nature 558, 440–444. doi:10.1038/s41586-018-0207-y

Dahal, B., NandaKafle, G., Perkins, L., Brözel, V.S., 2017. Diversity of free-Living nitrogen fixing Streptomyces in soils of the badlands of South Dakota. Microbiological Research 195, 31–39. doi:10.1016/j.micres.2016.11.004

Davidson, N.C., Fluet-Chouinard, E., Finlayson, C.M., 2018. Global extent and distribution of wetlands: trends and issues. Marine and Freshwater Research 69, 620. doi:10.1071/MF17019

Delmont, T.O., Quince, C., Shaiber, A., Esen, Ö.C., Lee, S.T., Rappé, M.S., McLellan, S.L., Lücker, S., Eren, A.M., 2018. Nitrogen-fixing populations of Planctomycetes and Proteobacteria are abundant in surface ocean metagenomes. Nature Microbiology 3, 804–813. doi:10.1038/s41564-018-0176-9

Dos Santos, P.C., Fang, Z., Mason, S.W., Setubal, J.C., Dixon, R., 2012. Distribution of nitrogen fixation and nitrogenase-like sequences amongst microbial genomes. BMC Genomics 13, 162. doi:10.1186/1471-2164-13-162

Drake, J.B., Weishampel, J.F., 2000. Multifractal analysis of canopy height measures in a longleaf pine savanna. Forest Ecology and Management 128, 121–127. doi:10.1016/S0378-1127(99)00279-0

Eddy, S.R., 2011. Accelerated Profile HMM Searches. PLoS Computational Biology 7, e1002195. doi:10.1371/journal.pcbi.1002195

Edgar, R.C., 2016. SINTAX: a simple non-Bayesian taxonomy classifier for 16S and ITS sequences. BioRxiv. doi:http://doi.org/10.1101/074161

Edgar, R.C., 2010. Search and clustering orders of magnitude faster than BLAST. Bioinformatics 26, 2460–2461. doi:10.1093/bioinformatics/btq461

Fan, K., Delgado-Baquerizo, M., Guo, X., Wang, D., Wu, Y., Zhu, M., Yu, W., Yao, H., Zhu, Y., Chu, H., 2019. Suppressed N fixation and diazotrophs after four decades of fertilization. Microbiome 7, 143. doi:10.1186/s40168-019-0757-8

Federhen, S., 2012. The NCBI Taxonomy database. Nucleic Acids Research 40, D136–D143. doi:10.1093/nar/gkr1178

Fodelianakis, S., Valenzuela-Cuevas, A., Barozzi, A., Daffonchio, D., 2021. Direct quantification of ecological drift at the population level in synthetic bacterial communities. The ISME Journal 15, 55–66. doi:10.1038/s41396-020-00754-4

Gaby, J.C., Buckley, D.H., 2011. A global census of nitrogenase diversity. Environmental Microbiology 13, 1790–1799. doi:10.1111/j.1462-2920.2011.02488.x

Hartman, W.H., Ye, R., Horwath, W.R., Tringe, S.G., 2017. A genomic perspective on stoichiometric regulation of soil carbon cycling. The ISME Journal 11, 2652–2665. doi:10.1038/ismej.2017.115

Hsu, S.-F., Buckley, D.H., 2009. Evidence for the functional significance of diazotroph community structure in soil. The ISME Journal 3, 124–136. doi:10.1038/ismej.2008.82

Huber, D.H., Ugwuanyi, I.R., Malkaram, S.A., Montenegro-Garcia, N.A., Lhilhi Noundou, V., Chavarria-Palma, J.E., 2018. Metagenome Sequences of Sediment from a Recovering Industrialized Appalachian River in West Virginia. Genome Announcements 6. doi:10.1128/genomeA.00350-18

Hug, L.A., Baker, B.J., Anantharaman, K., Brown, C.T., Probst, A.J., Castelle, C.J., Butterfield, C.N., Hernsdorf, A.W., Amano, Y., Ise, K., Suzuki, Y., Dudek, N., Relman, D.A., Finstad, K.M., Amundson, R., Thomas, B.C., Banfield, J.F., 2016. A new view of the tree of life. Nature Microbiology 1, 16048. doi:10.1038/nmicrobiol.2016.48

Hyatt, D., Chen, G.-L., LoCascio, P.F., Land, M.L., Larimer, F.W., Hauser, L.J., 2010. Prodigal: prokaryotic gene recognition and translation initiation site identification. BMC Bioinformatics 11, 119. doi:10.1186/1471-2105-11-119

Itoh, H., Xu, Z., Masuda, Y., Ushijima, N., Hayakawa, C., Shiratori, Y., Senoo, K., 2021. Geomonas silvestris sp. nov., Geomonas paludis sp. nov. and Geomonas limicola sp. nov., isolated from terrestrial environments, and emended description of the genus Geomonas. International Journal of Systematic and Evolutionary Microbiology 71, 004607. doi:10.1099/ijsem.0.004607

Itoh, H., Xu, Z., Mise, K., Masuda, Y., Ushijima, N., Hayakawa, C., Shiratori, Y., Senoo, K., 2022. Anaeromyxobacter oryzae sp. nov., Anaeromyxobacter diazotrophicus sp. nov. and Anaeromyxobacter paludicola sp. nov., isolated from paddy soils. International Journal of Systematic and Evolutionary Microbiology 72, 005546. doi:10.1099/ijsem.0.005546

Jiang, H., Zhou, R., Zhang, M., Cheng, Z., Li, J., Zhang, G., Chen, B., Zou, S., Yang, Y., 2018. Exploring the differences of antibiotic resistance genes profiles between river surface water and sediments using metagenomic approach. Ecotoxicology and Environmental Safety 161, 64–69. doi:10.1016/j.ecoenv.2018.05.044

Johnston, E.R., Rodriguez-R, L.M., Luo, C., Yuan, M.M., Wu, L., He, Z., Schuur, E.A.G., Luo, Y., Tiedje, J.M., Zhou, J., Konstantinidis, K.T., 2016. Metagenomics Reveals Pervasive Bacterial Populations and Reduced Community Diversity across the Alaska Tundra Ecosystem. Frontiers in Microbiology 7, 579. doi:10.3389/fmicb.2016.00579

Jones, C.M., Graf, D.R., Bru, D., Philippot, L., Hallin, S., 2013. The unaccounted yet abundant nitrous oxide-reducing microbial community: a potential nitrous oxide sink. The ISME Journal 7, 417–426. doi:10.1038/ismej.2012.125

Kanehisa, M., Furumichi, M., Sato, Y., Ishiguro-Watanabe, M., Tanabe, M., 2021. KEGG: Integrating viruses and cellular organisms. Nucleic Acids Research 49, D545–D551. doi:10.1093/nar/gkaa970

Kapili, B.J., Dekas, A.E., 2021. PPIT: an R package for inferring microbial taxonomy from nifH sequences. Bioinformatics 37, 2289–2298. doi:10.1093/bioinformatics/btab100

Katoh, K., 2002. MAFFT: a novel method for rapid multiple sequence alignment based on fast Fourier transform. Nucleic Acids Research 30, 3059–3066. doi:10.1093/nar/gkf436

Katz, K., Shutov, O., Lapoint, R., Kimelman, M., Brister, J.R., O’Sullivan, C., 2022. The Sequence Read Archive: a decade more of explosive growth. Nucleic Acids Research 50, D387–D390. doi:10.1093/nar/gkab1053

Kim, M., Oh, H.-S., Park, S.-C., Chun, J., 2014. Towards a taxonomic coherence between average nucleotide identity and 16S rRNA gene sequence similarity for species demarcation of prokaryotes. International Journal of Systematic and Evolutionary Microbiology 64, 346–351. doi:10.1099/ijs.0.059774-0

Kopylova, E., Noé, L., Touzet, H., 2012. SortMeRNA: fast and accurate filtering of ribosomal RNAs in metatranscriptomic data. Bioinformatics 28, 3211–3217. doi:10.1093/bioinformatics/bts611

Kuypers, M.M.M., Marchant, H.K., Kartal, B., 2018. The microbial nitrogen-cycling network. Nature Reviews Microbiology 16, 263–276. doi:10.1038/nrmicro.2018.9

Letunic, I., Bork, P., 2021. Interactive Tree Of Life (iTOL) v5: an online tool for phylogenetic tree display and annotation. Nucleic Acids Research 49, W293–W296. doi:10.1093/nar/gkab301

Li, H.-Y., Wang, H., Wang, H.-T., Xin, P.-Y., Xu, X.-H., Ma, Y., Liu, W.-P., Teng, C.-Y., Jiang, C.-L., Lou, L.-P., Arnold, W., Cralle, L., Zhu, Y.-G., Chu, J.-F., Gilbert, J.A., Zhang, Z.-J., 2018. The chemodiversity of paddy soil dissolved organic matter correlates with microbial community at continental scales. Microbiome 6, 187. doi:10.1186/s40168-018-0561-x

Li, Y., Tremblay, J., Bainard, L.D., Cade-Menun, B., Hamel, C., 2020. Long-term effects of nitrogen and phosphorus fertilization on soil microbial community structure and function under continuous wheat production. Environmental Microbiology 22, 1066–1088. doi:10.1111/1462-2920.14824

Links, M.G., Dumonceaux, T.J., McCarthy, E.L., Hemmingsen, S.M., Topp, E., Town, J.R., 2021. CaptureSeq: Hybridization-Based Enrichment of cpn60 Gene Fragments Reveals the Community Structures of Synthetic and Natural Microbial Ecosystems. Microorganisms 9, 816. doi:10.3390/microorganisms9040816

Liu, G.-H., Yang, S., Tang, R., Xie, C.-J., Zhou, S.-G., 2021. Genome Analysis and Description of Three Novel Diazotrophs Geomonas Species Isolated From Paddy Soils. Frontiers in Microbiology 12, 801462. doi:10.3389/fmicb.2021.801462

Liu, Y.-R., Johs, A., Bi, L., Lu, X., Hu, H.-W., Sun, D., He, J.-Z., Gu, B., 2018. Unraveling Microbial Communities Associated with Methylmercury Production in Paddy Soils. Environmental Science & Technology 52, 13110–13118. doi:10.1021/acs.est.8b03052

Lovley, D.R., Giovannoni, S.J., White, D.C., Champine, J.E., Phillips, E.J.P., Gorby, Y.A., Goodwin, S., 1993. Geobacter metallireducens gen. nov. sp. nov., a microorganism capable of coupling the complete oxidation of organic compounds to the reduction of iron and other metals. Archives of Microbiology 159, 336–344. doi:10.1007/BF00290916

Ma, B., Zhao, K., Lv, X., Su, W., Dai, Z., Gilbert, J.A., Brookes, P.C., Faust, K., Xu, J., 2018. Genetic correlation network prediction of forest soil microbial functional organization. The ISME Journal 12, 2492–2505. doi:10.1038/s41396-018-0232-8

Mahmud, K., Makaju, S., Ibrahim, R., Missaoui, A., 2020. Current Progress in Nitrogen Fixing Plants and Microbiome Research. Plants 9, 97. doi:10.3390/plants9010097

Masuda, Y., Itoh, H., Shiratori, Y., Isobe, K., Otsuka, S., Senoo, K., 2017. Predominant but previously-overlooked prokaryotic drivers of reductive nitrogen transformation in paddy soils, revealed by metatranscriptomics. Microbes and Environments 32, 180–183. doi:10.1264/jsme2.ME16179

Masuda, Y., Yamanaka, H., Xu, Z.-X., Shiratori, Y., Aono, T., Amachi, S., Senoo, K., Itoh, H., 2020. Diazotrophic Anaeromyxobacter Isolates from Soils. Applied and Environmental Microbiology 86, e00956–20. doi:10.1128/AEM.00956-20

Matsen, F.A., Kodner, R.B., Armbrust, E.V., 2010. pplacer: linear time maximum-likelihood and Bayesian phylogenetic placement of sequences onto a fixed reference tree. BMC Bioinformatics 11, 538. doi:10.1186/1471-2105-11-538

Meyer, F., Bagchi, S., Chaterji, S., Gerlach, W., Grama, A., Harrison, T., Paczian, T., Trimble, W.L., Wilke, A., 2019. MG-RAST version 4—lessons learned from a decade of low-budget ultra-high-throughput metagenome analysis. Briefings in Bioinformatics 20, 1151–1159. doi:10.1093/bib/bbx105

Meyer, F., Paarmann, D., D’Souza, M., Olson, R., Glass, E., Kubal, M., Paczian, T., Rodriguez, A., Stevens, R., Wilke, A., Wilkening, J., Edwards, R., 2008. The metagenomics RAST server – a public resource for the automatic phylogenetic and functional analysis of metagenomes. BMC Bioinformatics 9, 386. doi:10.1186/1471-2105-9-386

Mise, K., Masuda, Y., Senoo, K., Itoh, H., 2021. Undervalued Pseudo-nifH Sequences in Public Databases Distort Metagenomic Insights into Biological Nitrogen Fixers. mSphere 6, e00785–21. doi:10.1128/msphere.00785-21

Mitter, E.K., Germida, J.J., de Freitas, J.R., 2021. Impact of diesel and biodiesel contamination on soil microbial community activity and structure. Scientific Reports 11, 10856. doi:10.1038/s41598-021-89637-y

Neal, A.L., Hughes, D., Clark, I.M., Jansson, J.K., Hirsch, P.R., 2021. Microbiome Aggregated Traits and Assembly Are More Sensitive to Soil Management than Diversity. mSystems 6. doi:10.1128/mSystems.01056-20

Nelkner, J., Henke, C., Lin, T.W., Pätzold, W., Hassa, J., Jaenicke, S., Grosch, R., Pühler, A., Sczyrba, A., Schlüter, A., 2019. Effect of Long-Term Farming Practices on Agricultural Soil Microbiome Members Represented by Metagenomically Assembled Genomes (MAGs) and Their Predicted Plant-Beneficial Genes. Genes 10, 424. doi:10.3390/genes10060424

Nelson, M.B., Martiny, A.C., Martiny, J.B.H., 2016. Global biogeography of microbial nitrogen-cycling traits in soil. Proceedings of the National Academy of Sciences 113, 8033–8040. doi:10.1073/pnas.1601070113

Nemergut, D.R., Schmidt, S.K., Fukami, T., O’Neill, S.P., Bilinski, T.M., Stanish, L.F., Knelman, J.E., Darcy, J.L., Lynch, R.C., Wickey, P., Ferrenberg, S., 2013. Patterns and Processes of Microbial Community Assembly. Microbiology and Molecular Biology Reviews 77, 342–356. doi:10.1128/MMBR.00051-12

Oksanen, J., Blanchet, F.G., Friendly, M., Kindt, R., Legendre, P., McGlinn, D., Minchin, P.R., O’Hara, R.B., Simpson, G.L., Solymos, P., Stevens, M.H.H., Szoecs, E., Wagner, H., 2020. vegan: Community Ecology Package.

Orellana, L.H., Chee-Sanford, J.C., Sanford, R.A., Löffler, F.E., Konstantinidis, K.T., 2018. Year-Round Shotgun Metagenomes Reveal Stable Microbial Communities in Agricultural Soils and Novel Ammonia Oxidizers Responding to Fertilization. Applied and Environmental Microbiology 84. doi:10.1128/AEM.01646-17

Ouyang, Y., Norton, J.M., 2020. Short-Term Nitrogen Fertilization Affects Microbial Community Composition and Nitrogen Mineralization Functions in an Agricultural Soil. Applied and Environmental Microbiology 86, 516–8. doi:10.1128/AEM.02278-19

Parks, D.H., Chuvochina, M., Rinke, C., Mussig, A.J., Chaumeil, P.-A., Hugenholtz, P., 2022. GTDB: an ongoing census of bacterial and archaeal diversity through a phylogenetically consistent, rank normalized and complete genome-based taxonomy. Nucleic Acids Research 50, D785–D794. doi:10.1093/nar/gkab776

Paungfoo-Lonhienne, C., Wang, W., Yeoh, Y.K., Halpin, N., 2017. Legume crop rotation suppressed nitrifying microbial community in a sugarcane cropping soil. Scientific Reports 7, 16707. doi:10.1038/s41598-017-17080-z

Pecher, W.T., Martínez, F.L., DasSarma, P., Guzmán, D., DasSarma, S., 2020. 16S rRNA Gene Diversity in Ancient Gray and Pink Salt from San Simón Salt Mines in Tarija, Bolivia. Microbiology Resource Announcements 9. doi:10.1128/MRA.00820-20

Price, M.N., Dehal, P.S., Arkin, A.P., 2010. FastTree 2 – Approximately Maximum-Likelihood Trees for Large Alignments. PLoS ONE 5, e9490. doi:10.1371/journal.pone.0009490

Price, M.N., Dehal, P.S., Arkin, A.P., 2009. FastTree: Computing Large Minimum Evolution Trees with Profiles instead of a Distance Matrix. Molecular Biology and Evolution 26, 1641–1650. doi:10.1093/molbev/msp077

Quast, C., Pruesse, E., Yilmaz, P., Gerken, J., Schweer, T., Yarza, P., Peplies, J., Glöckner, F.O., 2012. The SILVA ribosomal RNA gene database project: improved data processing and web-based tools. Nucleic Acids Research 41, D590–D596. doi:10.1093/nar/gks1219

R Core Team, 2021. R: A Language and Environment for Statistical Computing.

Robson, R.L., Postgate, J.R., 1980. Oxygen and Hydrogen in Biological Nitrogen Fixation. Annual Review of Microbiology 34, 183–207. doi:10.1146/annurev.mi.34.100180.001151

Romanowicz, K.J., Crump, B.C., Kling, G.W., 2021. Rainfall Alters Permafrost Soil Redox Conditions, but Meta-Omics Show Divergent Microbial Community Responses by Tundra Type in the Arctic. Soil Systems 5, 17. doi:10.3390/soilsystems5010017

Sevim, V., Lee, Juna, Egan, R., Clum, A., Hundley, H., Lee, Janey, Everroad, R.C., Detweiler, A.M., Bebout, B.M., Pett-Ridge, J., Göker, M., Murray, A.E., Lindemann, S.R., Klenk, H.-P., O’Malley, R., Zane, M., Cheng, J.-F., Copeland, A., Daum, C., Singer, E., Woyke, T., 2019. Shotgun metagenome data of a defined mock community using Oxford Nanopore, PacBio and Illumina technologies. Scientific Data 6, 285. doi:10.1038/s41597-019-0287-z

Shen, W., Le, S., Li, Y., Hu, F., 2016. SeqKit: A Cross-Platform and Ultrafast Toolkit for FASTA/Q File Manipulation. PLOS ONE 11, e0163962. doi:10.1371/journal.pone.0163962

Shen, W., Ren, H., 2021. TaxonKit: A practical and efficient NCBI taxonomy toolkit. Journal of Genetics and Genomics 48, 844–850. doi:10.1016/j.jgg.2021.03.006

Soumare, A., Diedhiou, A.G., Thuita, M., Hafidi, M., Ouhdouch, Y., Gopalakrishnan, S., Kouisni, L., 2020. Exploiting Biological Nitrogen Fixation: A Route Towards a Sustainable Agriculture. Plants 9, 1011. doi:10.3390/plants9081011

Sukhum, K. V., Vargas, R.C., Boolchandani, M., D’Souza, A.W., Patel, S., Kesaraju, A., Walljasper, G., Hegde, H., Ye, Z., Valenzuela, R.K., Gunderson, P., Bendixsen, C., Dantas, G., Shukla, S.K., 2021. Manure Microbial Communities and Resistance Profiles Reconfigure after Transition to Manure Pits and Differ from Those in Fertilized Field Soil. mBio 12. doi:10.1128/mBio.00798-21

Sun, W., Xiao, E., Pu, Z., Krumins, V., Dong, Y., Li, B., Hu, M., 2018. Paddy soil microbial communities driven by environment- and microbe-microbe interactions: A case study of elevation-resolved microbial communities in a rice terrace. Science of The Total Environment 612, 884–893. doi:10.1016/j.scitotenv.2017.08.275

Suttner, B., Johnston, E.R., Orellana, L.H., Rodriguez-R, L.M., Hatt, J.K., Carychao, D., Carter, M.Q., Cooley, M.B., Konstantinidis, K.T., 2020. Metagenomics as a Public Health Risk Assessment Tool in a Study of Natural Creek Sediments Influenced by Agricultural and Livestock Runoff: Potential and Limitations. Applied and Environmental Microbiology 86. doi:10.1128/AEM.02525-19

Takada-Hoshino, Y., Matsumoto, N., 2004. An Improved DNA Extraction Method Using Skim Milk from Soils That Strongly Adsorb DNA. Microbes and Environments 19, 13–19. doi:10.1264/jsme2.19.13

Vellend, M., 2010. Conceptual Synthesis in Community Ecology. The Quarterly Review of Biology 85, 183–206. doi:10.1086/652373

Vitousek, P.M., Cassman, K., Cleveland, C., Crews, T., Field, C.B., Grimm, N.B., Howarth, R.W., Marino, R., Martinelli, L., Rastetter, E.B., Sprent, J.I., 2002. Towards an ecological understanding of biological nitrogen fixation. Biogeochemistry 57, 1–45. doi:10.1023/A:1015798428743

Wang, Hang, He, X., Zhang, Z., Li, M., Zhang, Q., Zhu, H., Xu, S., Yang, P., 2021. Eight years of manure fertilization favor copiotrophic traits in paddy soil microbiomes. European Journal of Soil Biology 106, 103352. doi:10.1016/j.ejsobi.2021.103352

Wang, Huanhuan, Li, Xu, Li, Xinyu, Li, F., Su, Z., Zhang, H., 2021. Community Composition and Co-Occurrence Patterns of Diazotrophs along a Soil Profile in Paddy Fields of Three Soil Types in China. Microbial Ecology 82, 961–970. doi:10.1007/s00248-021-01716-9

Wang, J., Long, Z., Min, W., Hou, Z., 2020. Metagenomic analysis reveals the effects of cotton straw– derived biochar on soil nitrogen transformation in drip-irrigated cotton field. Environmental Science and Pollution Research 27, 43929–43941. doi:10.1007/s11356-020-10267-4

Wang, Q., Quensen, J.F., Fish, J.A., Kwon Lee, T., Sun, Y., Tiedje, J.M., Cole, J.R., 2013. Ecological Patterns of nifH Genes in Four Terrestrial Climatic Zones Explored with Targeted Metagenomics Using FrameBot, a New Informatics Tool. mBio 4. doi:10.1128/mBio.00592-13

Wang, X., Teng, Y., Ren, W., Li, Y., Yang, T., Chen, Y., Zhao, L., Zhang, H., Kuramae, E.E., 2022. Variations of Bacterial and Diazotrophic Community Assemblies throughout the Soil Profile in Distinct Paddy Soil Types and Their Contributions to Soil Functionality. mSystems. doi:10.1128/msystems.01047-21

Weber, K.A., Achenbach, L.A., Coates, J.D., 2006. Microorganisms pumping iron: anaerobic microbial iron oxidation and reduction. Nature Reviews Microbiology 4, 752–764. doi:10.1038/nrmicro1490

Woodcroft, B.J., Singleton, C.M., Boyd, J.A., Evans, P.N., Emerson, J.B., Zayed, A.A.F., Hoelzle, R.D., Lamberton, T.O., McCalley, C.K., Hodgkins, S.B., Wilson, R.M., Purvine, S.O., Nicora, C.D., Li, C., Frolking, S., Chanton, J.P., Crill, P.M., Saleska, S.R., Rich, V.I., Tyson, G.W., 2018. Genome-centric view of carbon processing in thawing permafrost. Nature 560, 49–54. doi:10.1038/s41586-018-0338-1

Wu, D., Zhao, Y., Cheng, L., Zhou, Z., Wu, Q., Wang, Q., Yuan, Q., 2021. Activity and structure of methanogenic microbial communities in sediments of cascade hydropower reservoirs, Southwest China. Science of The Total Environment 786, 147515. doi:10.1016/j.scitotenv.2021.147515

Xiao, K.-Q., Li, B., Ma, L., Bao, P., Zhou, X., Zhang, T., Zhu, Y.-G., 2016. Metagenomic profiles of antibiotic resistance genes in paddy soils from South China. FEMS Microbiology Ecology 92, fiw023. doi:10.1093/femsec/fiw023

Xu, X., Thornton, P.E., Post, W.M., 2013. A global analysis of soil microbial biomass carbon, nitrogen and phosphorus in terrestrial ecosystems. Global Ecology and Biogeography 22, 737–749. doi:10.1111/geb.12029

Xu, Z., Masuda, Y., Hayakawa, C., Ushijima, N., Kawano, K., Shiratori, Y., Senoo, K., Itoh, H., 2020. Description of Three Novel Members in the Family Geobacteraceae, Oryzomonas japonicum gen. nov., sp. nov., Oryzomonas sagensis sp. nov., and Oryzomonas ruber sp. nov. Microorganisms 8, 634. doi:10.3390/microorganisms8050634

Xu, Z., Masuda, Y., Itoh, H., Ushijima, N., Shiratori, Y., Senoo, K., 2019. Geomonas oryzae gen. nov., sp. nov., Geomonas edaphica sp. nov., Geomonas ferrireducens sp. nov., Geomonas terrae sp. nov., Four Ferric-Reducing Bacteria Isolated From Paddy Soil, and Reclassification of Three Species of the Genus Geobacter as Members of the Genus Geomonas gen. nov. Frontiers in Microbiology 10, 2201. doi:10.3389/fmicb.2019.02201

Xu, Z., Masuda, Y., Wang, X., Ushijima, N., Shiratori, Y., Senoo, K., Itoh, H., 2021. Genome-Based Taxonomic Rearrangement of the Order Geobacterales Including the Description of Geomonas azotofigens sp. nov. and Geomonas diazotrophica sp. nov. Frontiers in Microbiology 12, 2715. doi:10.3389/fmicb.2021.737531

Xue, Y., Jonassen, I., Øvreås, L., Taş, N., 2019. Bacterial and Archaeal Metagenome-Assembled Genome Sequences from Svalbard Permafrost. Microbiology Resource Announcements 8. doi:10.1128/MRA.00516-19

Yang, S., Liu, G.-H., Tang, R., Han, S., Xie, C.-J., Zhou, S.-G., 2022. Description of two nitrogen-fixing bacteria, Geomonas fuzhouensis sp. nov. and Geomonas agri sp. nov., isolated from paddy soils. Antonie van Leeuwenhoek 115, 435–444. doi:10.1007/s10482-021-01704-6

Yu, J., Deem, L.M., Crow, S.E., Deenik, J., Penton, C.R., 2019. Comparative Metagenomics Reveals Enhanced Nutrient Cycling Potential after 2 Years of Biochar Amendment in a Tropical Oxisol. Applied and Environmental Microbiology 85. doi:10.1128/AEM.02957-18

Yu, K., Zhang, T., 2013. Construction of Customized Sub-Databases from NCBI-nr Database for Rapid Annotation of Huge Metagenomic Datasets Using a Combined BLAST and MEGAN Approach. PLoS ONE 8, e59831. doi:10.1371/journal.pone.0059831

Yu, Y., Zhang, J., Petropoulos, E., Baluja, M.Q., Zhu, C., Zhu, J., Lin, X., Feng, Y., 2018. Divergent Responses of the Diazotrophic Microbiome to Elevated CO_2_in Two Rice Cultivars. Frontiers in Microbiology 9. doi:10.3389/fmicb.2018.01139

Yurgel, S.N., Nearing, J.T., Douglas, G.M., Langille, M.G.I., 2019. Metagenomic Functional Shifts to Plant Induced Environmental Changes. Frontiers in Microbiology 10, 1682. doi:10.3389/fmicb.2019.01682

Zhalnina, K., Louie, K.B., Hao, Z., Mansoori, N., da Rocha, U.N., Shi, S., Cho, H., Karaoz, U., Loqué, D., Bowen, B.P., Firestone, M.K., Northen, T.R., Brodie, E.L., 2018. Dynamic root exudate chemistry and microbial substrate preferences drive patterns in rhizosphere microbial community assembly. Nature Microbiology 3, 470–480. doi:10.1038/s41564-018-0129-3

Zhang, C., Song, Z., Zhuang, D., Wang, J., Xie, S., Liu, G., 2019. Urea fertilization decreases soil bacterial diversity, but improves microbial biomass, respiration, and N-cycling potential in a semiarid grassland. Biology and Fertility of Soils 55, 229–242. doi:10.1007/s00374-019-01344-z

Zhang, Z., Xu, Z., Masuda, Y., Wang, X., Ushijima, N., Shiratori, Y., Senoo, K., Itoh, H., 2021. Geomesophilobacter sediminis gen. nov., sp. nov., Geomonas propionica sp. nov. and Geomonas anaerohicana sp. nov., three novel members in the family Geobacterecace isolated from river sediment and paddy soil. Systematic and Applied Microbiology 44, 126233. doi:10.1016/j.syapm.2021.126233

Zheng, Z., Li, L., Makhalanyane, T.P., Xu, Chunming, Li, K., Xue, K., Xu, Cong, Qian, R., Zhang, B., Du, J., Yu, H., Cui, X., Wang, Y., Hao, Y., 2021. The composition of antibiotic resistance genes is not affected by grazing but is determined by microorganisms in grassland soils. Science of The Total Environment 761, 143205. doi:10.1016/j.scitotenv.2020.143205

Zhu, C., Friman, V., Li, L., Xu, Q., Guo, J., Guo, S., Shen, Q., Ling, N., 2022. Meta-analysis of diazotrophic signatures across terrestrial ecosystems at the continental scale. Environmental Microbiology 24, 2013–2028. doi:10.1111/1462-2920.15984

