## Supplementary material for "Global soil metagenomics reveals ubiquitous yet previously-hidden predominance of *Deltaproteobacteria* in nitrogen-fixing microbiome": Figs S1-S3

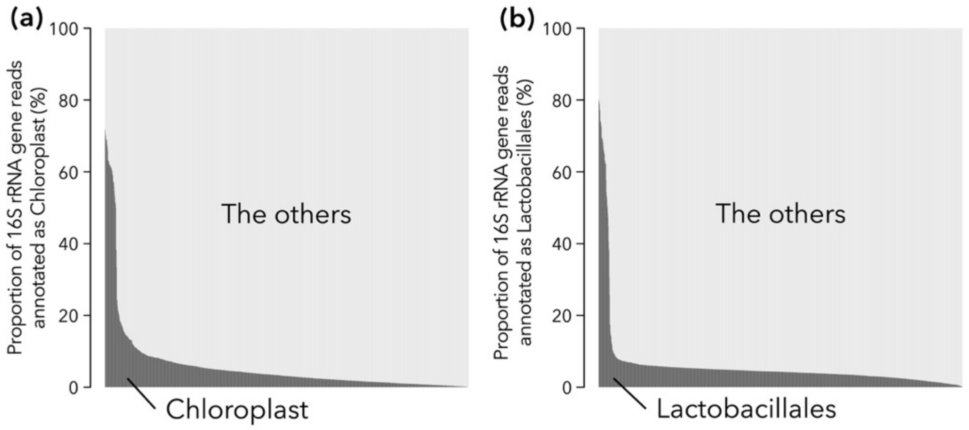


**Figure S1**. Proportion of 16S rRNA gene reads annotated as chloroplast (left panel) or order *Lactobacillales* (right panel) in each of the 1,445 metagenomes, presented in descending order.

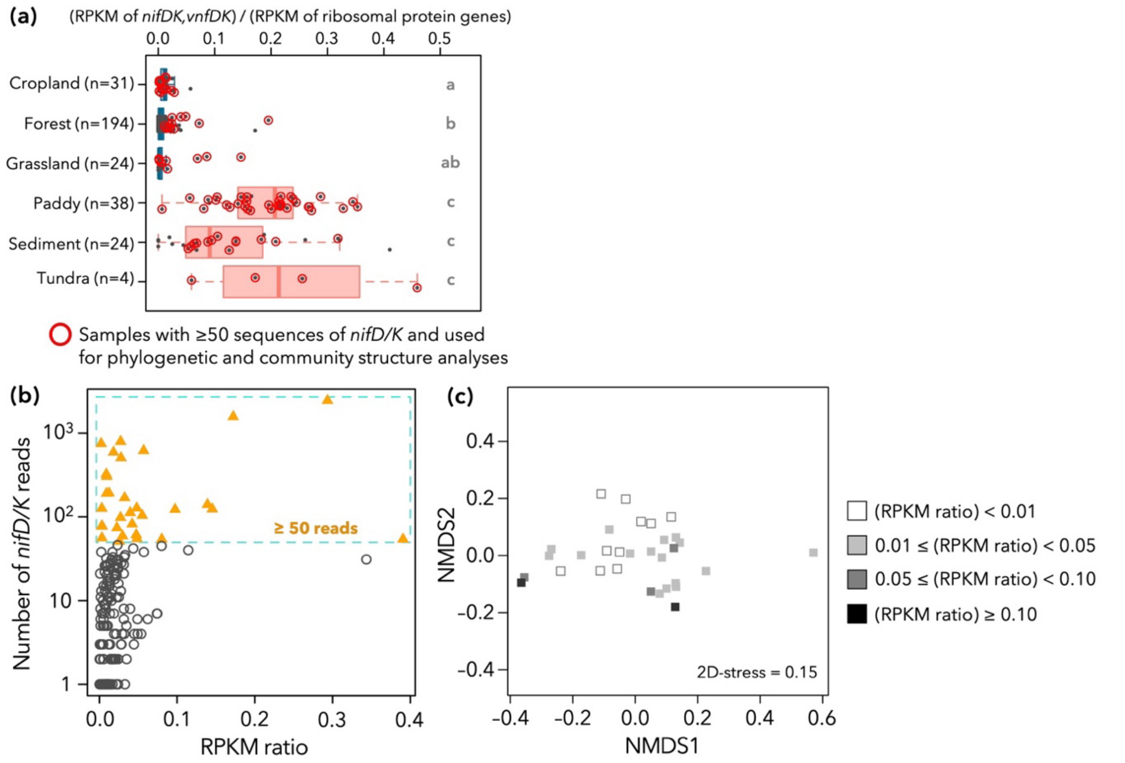


**Figure S2**. Potential bias introduced by using only datasets with 50 or more sequences of *nifD/K*. (a) The dominance of nitrogen-fixing prokaryotes in each environment. The figure is identical to Fig.1d, with samples containing 50 or more sequences of *nifD/K* (i.e., used for community structure analyses) are highlighted in red circles. (b) The relationship between RPKM ratio of nitrogenase to ribosomal protein genes and the number of *nifD/K* reads. Each dot represents one dataset. Datasets containing 50 or more sequences of *nifD/K* (i.e., used for community structure analyses) are depicted in pink and highlighted by a red dotted square. (c) The overall beta-diversity of *nifD* and *nifK* sequences summarized by nonmetric multidimensional scaling (NMDS) in cropland, forest, and grassland datasets. Symbols are colored according to the dominance of nitrogen-fixing prokaryotes (represented by RPKM ratios).

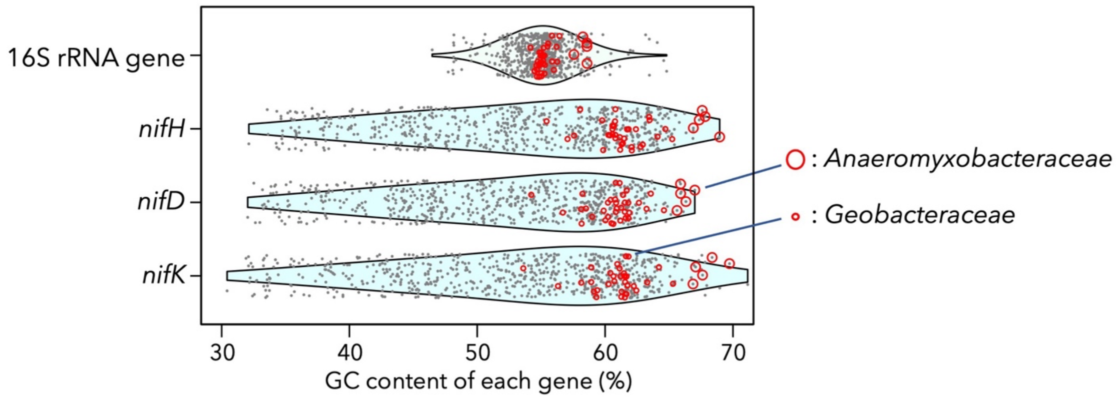


**Figure S3**. GC contents in 16S rRNA genes and nitrogenase genes (*nifH/D/K*) on prokaryotic genomes. Each point corresponds to one genome, and those denoting *Anaeromyxobacteraceae* and *Geobacteraceae* genomes are indicated by red circles (large and small ones, respectively). Violin plots indicate the density of points.
